## Supplementary figures and images for "Fifteenth century CE Bolivian maize reveals genetic affinities with ancient Peruvian maize"

### Supplemental Figure 1

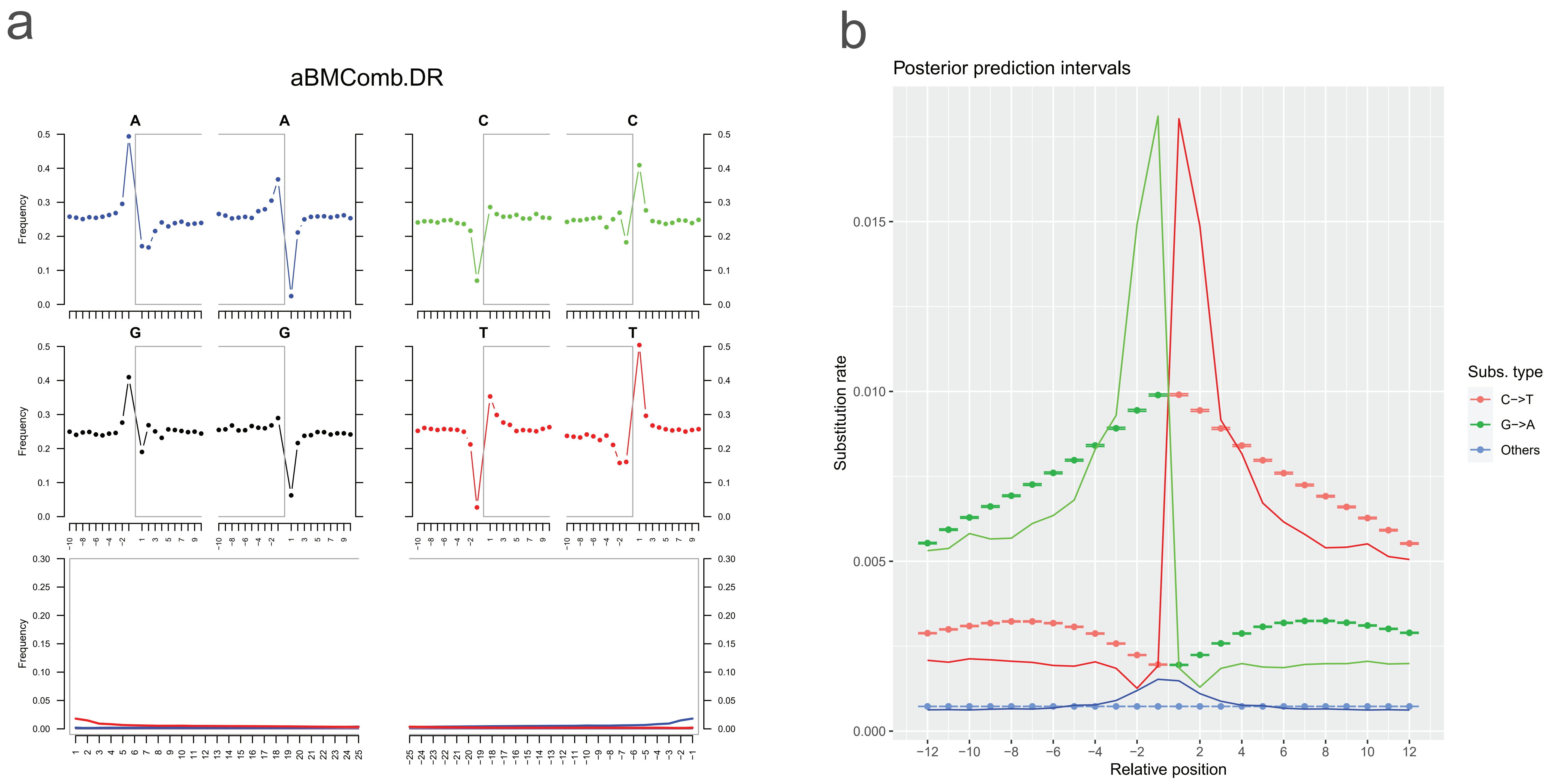

### Supplemental Figure 2

#### aBMComb.DR.srt

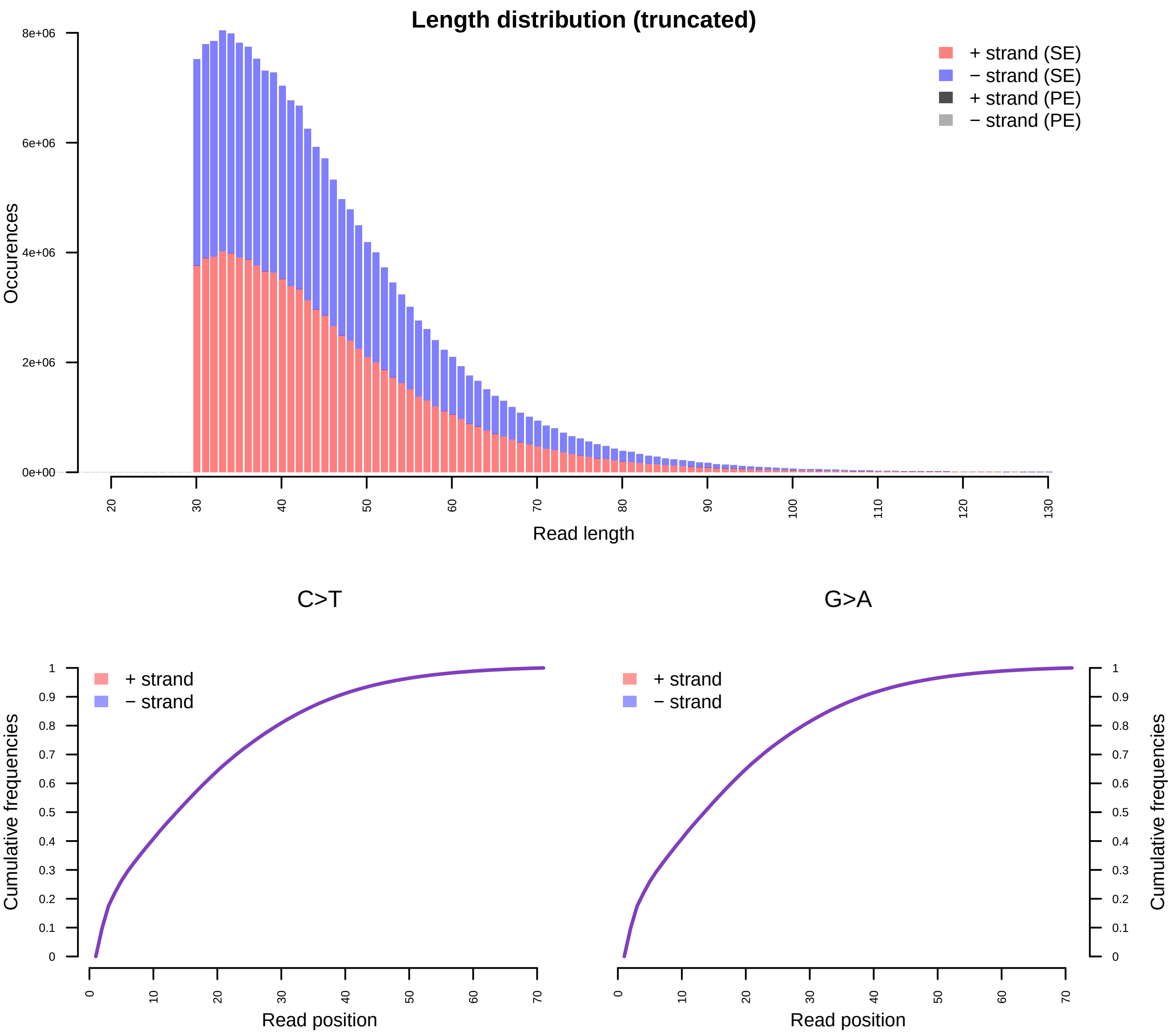

### Supplemental Figure 3

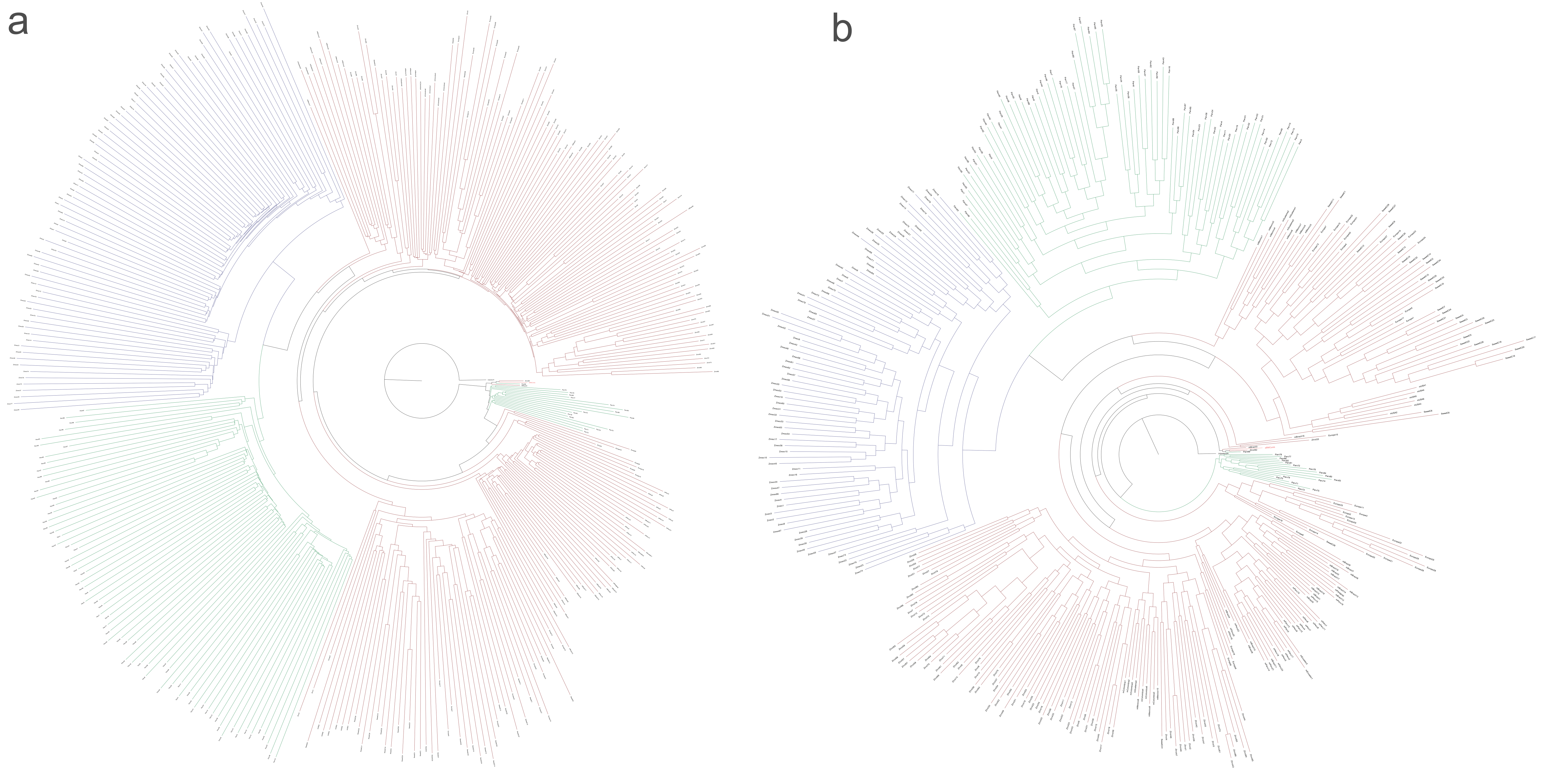

### Supplemental Figure 4

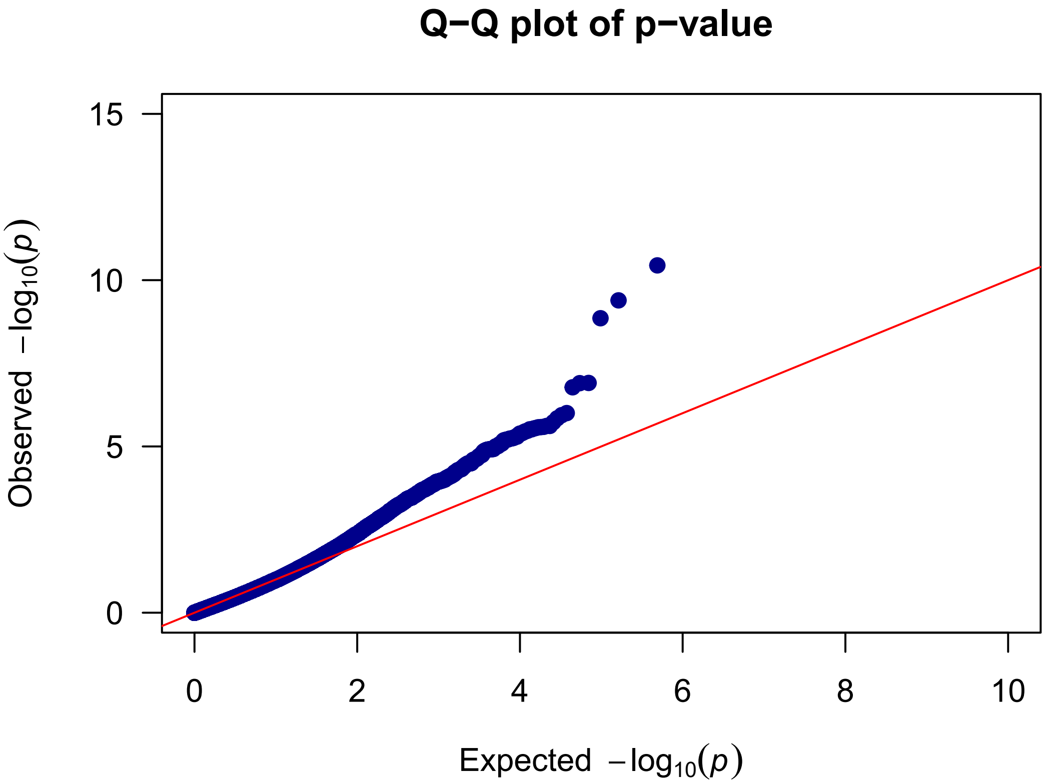
