## Supplementa1 Dataset S1 for "Fifteenth century CE Bolivian maize reveals genetic affinities with ancient Peruvian maize"

**Supplementary Dataset 1.** Modern Sample information.

| Number | ID | IDname | Publication | Grzybowski et al.2023_ID | Organism |
|---|---|---|---|---|---|
| 1 | USA_HotevillaArizona | mUSA1 | Wang et al. 2017 | RIMMA0415 | Zea mays |
| 2 | USA_AcomaPueblo | mUSA2 | Wang et al. 2017 | RIMMA0383 | Zea mays |
| 3 | USA_JemezPueblo | mUSA3 | Wang et al. 2017 | RIMMA0387 | Zea mays |
| 4 | USA_SanLorenzoPueblo | mUSA4 | Wang et al. 2017 | RIMMA0384 | Zea mays |
| 5 | USA_TesuquePueblo | mUSA5 | Wang et al. 2017 | RIMMA1012 | Zea mays |
| 6 | USA_TaosPueblo | mUSA6 | Wang et al. 2017 | RIMMA0385 | Zea mays |
| 7 | Mexico_Jalisco | mMexico1 | Wang et al. 2017 | RIMMA0623 | Zea mays |
| 8 | Mexico_Zacatecas | mMexico2 | Wang et al. 2017 | RIMMA0677 | Zea mays |
| 9 | Mexico | mMexico3 | Wang et al. 2017 | RIMMA0672 | Zea mays |
| 10 | Mexico_LaConcordiaGuerrero | mMexico4 | Wang et al. 2017 | RIMMA1010 | Zea mays |
| 11 | Mexico_Puebla1 | mMexico5 | Wang et al. 2017 | RIMMA0421 | Zea mays |
| 12 | Mexico_Puebla2 | mMexico6 | Wang et al. 2017 | RIMMA0626 | Zea mays |
| 13 | Mexico_Puebla3 | mMexico7 | Wang et al. 2017 | RIMMA0625 | Zea mays |
| 14 | Mexico_Oaxac | mMexico8 | Wang et al. 2017 | RIMMA0733 | Zea mays |
| 15 | Mexico_Chiapas | mMexico9 | Wang et al. 2017 | RIMMA0409 | Zea mays |
| 16 | Guatemala_SanMarcos1 | mGuatema1 | Wang et al. 2017 | RIMMA1007 | Zea mays |
| 17 | Guatemala_SanMarcos2 | mGuatema2 | Wang et al. 2017 | RIMMA0670 | Zea mays |
| 18 | Guatemala_Totonicapan | mGuatema3 | Wang et al. 2017 | RIMMA1008 | Zea mays |
| 19 | Guatemala | mGuatema4 | Wang et al. 2017 | RIMMA0720 | Zea mays |
| 20 | Mexico_Yucatan | mMexico10 | Wang et al. 2017 | RIMMA0703 | Zea mays |
| 21 | Ecuador1 | mEcuador1 | Wang et al. 2017 | RIMMA0665 | Zea mays |
| 22 | Ecuador2 | mEcuador2 | Wang et al. 2017 | RIMMA0662 | Zea mays |
| 23 | Peru_Ancash1 | mPeru13 | Wang et al. 2017 | RIMMA0438 | Zea mays |
| 24 | Colombia_Choco | mColombia1 | Wang et al. 2017 | RIMMA0395 | Zea mays |
| 25 | Peru_Ancash2 | mPeru17 | Wang et al. 2017 | RIMMA0468 | Zea mays |
| 26 | Colombia_Magdalena1 | mColombia2 | Wang et al. 2017 | RIMMA0398 | Zea mays |
| 27 | Colombia_Caldas | mColombia3 | Wang et al. 2017 | RIMMA0390 | Zea mays |
| 28 | Colombia_Caqueta | mColombia4 | Wang et al. 2017 | RIMMA0392 | Zea mays |
| 29 | Colombia_Cordoba | mColombia5 | Wang et al. 2017 | RIMMA0393 | Zea mays |
| 30 | Colombia_Magdalena2 | mColombia6 | Wang et al. 2017 | RIMMA0399 | Zea mays |
| 31 | Peru_Apurimac | mPeru3 | Wang et al. 2017 | RIMMA0466 | Zea mays |

| | | | | | |
|---|---|---|---|---|---|
| 32 | Peru_Lalibertad1 | mPeru4 | Kistler et al. 2018 | C1_Kistler | Zea mays |
| 33 | Peru_Lima | mPeru5 | Kistler et al. 2018 | C10_Kistler | Zea mays |
| 34 | Peru_Apurimac1 | mPeru6 | Kistler et al. 2018 | C11_Kistler | Zea mays |
| 35 | Peru_Ica | mPeru7 | Kistler et al. 2018 | C12_Kistler | Zea mays |
| 36 | Peru_Arequipa | mPeru8 | Kistler et al. 2018 | C13_Kistler | Zea mays |
| 37 | Peru_Lambayeque | mPeru9 | Kistler et al. 2018 | C14_Kistler | Zea mays |
| 38 | Peru_Ayacucho | mPeru10 | Kistler et al. 2018 | C15_Kistler | Zea mays |
| 39 | Peru_Amazonas | mPeru11 | Kistler et al. 2018 | C17_Kistler | Zea mays |
| 40 | Peru_Lalibertad2 | mPeru12 | Kistler et al. 2018 | C18_Kistler | Zea mays |
| 41 | Peru_Ancash3 | mPeru1 | Kistler et al. 2018 | C19_Kistler | Zea mays |
| 42 | Peru_Apurimac2 | mPeru14 | Kistler et al. 2018 | C20_Kistler | Zea mays |
| 43 | Peru_SanMartin | mPeru15 | Kistler et al. 2018 | C21_Kistler | Zea mays |
| 44 | Peru_Apurimac3 | mPeru16 | Kistler et al. 2018 | C22_Kistler | Zea mays |
| 45 | Peru_Ancash4 | mPeru2 | Kistler et al. 2018 | C23_Kistler | Zea mays |
| 46 | Peru_Puno | mPeru18 | Kistler et al. 2018 | C26_Kistler | Zea mays |
| 47 | Brazil_Acre1 | mBrazil1 | Kistler et al. 2018 | C27_Kistler | Zea mays |
| 48 | Brazil_Rondonia1 | mBrazil2 | Kistler et al. 2018 | C28_Kistler | Zea mays |
| 49 | Brazil_Parana | mBrazil3 | Kistler et al. 2018 | C29_Kistler | Zea mays |
| 50 | Peru_Apurimac4 | mPeru19 | Kistler et al. 2018 | C3_Kistler | Zea mays |
| 51 | Paraguay | mParaguay | Kistler et al. 2018 | C30_Kistler | Zea mays |
| 52 | Brazil_Para1 | mBrazil4 | Kistler et al. 2018 | C31_Kistler | Zea mays |
| 53 | Brazil_SaoPaulo1 | mBrazil5 | Kistler et al. 2018 | C32_Kistler | Zea mays |
| 54 | Brazil_Para2 | mBrazil6 | Kistler et al. 2018 | C33_Kistler | Zea mays |
| 55 | Brazil_Altamira | mBrazil7 | Kistler et al. 2018 | C34_Kistler | Zea mays |
| 56 | Brazil_Maranhao | mBrazil8 | Kistler et al. 2018 | C35_Kistler | Zea mays |
| 57 | Brazil_Acre2 | mBrazil9 | Kistler et al. 2018 | C36_Kistler | Zea mays |
| 58 | Brazil_SaoPaulo2 | mBrazil10 | Kistler et al. 2018 | C38_Kistler | Zea mays |
| 59 | Brazil_Rondonia2 | mBrazil11 | Kistler et al. 2018 | C39_Kistler | Zea mays |
| 60 | Brazil_Acre3 | mBrazil12 | Kistler et al. 2018 | C40_Kistler | Zea mays |
| 61 | Brazil_MatoGrossoSul | mBrazil13 | Kistler et al. 2018 | C42_Kistler | Zea mays |
| 62 | Brazil_Para3 | mBrazil14 | Kistler et al. 2018 | C43_Kistler | Zea mays |
| 63 | Brazil_Rondonia3 | mBrazil15 | Kistler et al. 2018 | C44_Kistler | Zea mays |
| 64 | Brazil_MatoGrosso | mBrazil16 | Kistler et al. 2018 | C45_Kistler | Zea mays |

| | | | | | |
|---|---|---|---|---|---|
| 65 | Brazil_Goias | mBrazil17 | Kistler et al. 2018 | C46_Kistler | Zea mays |
| 66 | Brazil_Roraima4 | mBrazil18 | Kistler et al. 2018 | C47_Kistler | Zea mays |
| 67 | Brazil_Roraima5 | mBrazil19 | Kistler et al. 2018 | C48_Kistler | Zea mays |
| 68 | Peru_Junin | mPeru20 | Kistler et al. 2018 | C5_Kistler | Zea mays |
| 69 | Peru_Apurimac5 | mPeru21 | Kistler et al. 2018 | C6_Kistler | Zea mays |
| 70 | Peru_MadredeDios | mPeru22 | Kistler et al. 2018 | C8_Kistler | Zea mays |
| 71 | Peru_Apurimac6 | mPeru23 | Kistler et al. 2018 | C9_Kistler | Zea mays |
| 72 | Parviglumis_1 | Parviglumis1 | Chen et al., 2022 | 5A10 | Z. parviglumis |
| 73 | Parviglumis_2 | Parviglumis2 | Chen et al., 2022 | 5A11 | Z. parviglumis |
| 74 | Parviglumis_3 | Parviglumis3 | Chen et al., 2022 | 5A8 | Z. parviglumis |
| 75 | Parviglumis_4 | Parviglumis4 | Chen et al., 2022 | 5A9 | Z. parviglumis |
| 76 | Parviglumis_5 | Parviglumis5 | Chen et al., 2022 | 5B2 | Z. parviglumis |
| 77 | Parviglumis_6 | Parviglumis6 | Chen et al., 2022 | 5B3 | Z. parviglumis |
| 78 | Parviglumis_7 | Parviglumis7 | Chen et al., 2022 | 5B4 | Z. parviglumis |
| 79 | Parviglumis_8 | Parviglumis8 | Chen et al., 2022 | 5B5 | Z. parviglumis |
| 80 | Parviglumis_9 | Parviglumis9 | Chen et al., 2022 | 5B6 | Z. parviglumis |
| 81 | Parviglumis_10 | Parviglumis10 | Chen et al., 2022 | 5C1 | Z. parviglumis |
| 82 | Parviglumis_11 | Parviglumis11 | Chen et al., 2022 | 5C10 | Z. parviglumis |
| 83 | Parviglumis_12 | Parviglumis12 | Chen et al., 2022 | 5C11 | Z. parviglumis |
| 84 | Parviglumis_13 | Parviglumis13 | Chen et al., 2022 | 5C12 | Z. parviglumis |
| 85 | Parviglumis_14 | Parviglumis14 | Chen et al., 2022 | 5C2 | Z. parviglumis |
| 86 | Parviglumis_15 | Parviglumis15 | Chen et al., 2022 | 5C3 | Z. parviglumis |
| 87 | Parviglumis_16 | Parviglumis16 | Chen et al., 2022 | 5C4 | Z. parviglumis |
| 88 | Parviglumis_17 | Parviglumis17 | Chen et al., 2022 | 5C5 | Z. parviglumis |
| 89 | Parviglumis_18 | Parviglumis18 | Chen et al., 2022 | 5C6 | Z. parviglumis |
| 90 | Parviglumis_19 | Parviglumis19 | Chen et al., 2022 | 5C7 | Z. parviglumis |
| 91 | Parviglumis_20 | Parviglumis20 | Chen et al., 2022 | 5C8 | Z. parviglumis |
| 92 | Parviglumis_21 | Parviglumis21 | Chen et al., 2022 | 5C9 | Z. parviglumis |
| 93 | Parviglumis_22 | Parviglumis22 | Chen et al., 2022 | 5D10 | Z. parviglumis |
| 94 | Parviglumis_23 | Parviglumis23 | Chen et al., 2022 | 5D11 | Z. parviglumis |
| 95 | Parviglumis_24 | Parviglumis24 | Chen et al., 2022 | 5D12 | Z. parviglumis |
| 96 | Parviglumis_25 | Parviglumis25 | Chen et al., 2022 | 5D4 | Z. parviglumis |
| 97 | Parviglumis_26 | Parviglumis26 | Chen et al., 2022 | 5E1 | Z. parviglumis |

| | | | | | |
|---|---|---|---|---|---|
| 98 | Parviglumis_27 | Parviglumis27 | Chen et al., 2022 | 5E2 | Z. parviglumis |
| 99 | Parviglumis_28 | Parviglumis28 | Chen et al., 2022 | 5E3 | Z. parviglumis |
| 100 | Parviglumis_29 | Parviglumis29 | Chen et al., 2022 | 5E5 | Z. parviglumis |
| 101 | Parviglumis_30 | Parviglumis30 | Chen et al., 2022 | 5E6 | Z. parviglumis |
| 102 | Parviglumis_31 | Parviglumis31 | Chen et al., 2022 | 5F11 | Z. parviglumis |
| 103 | Parviglumis_32 | Parviglumis32 | Chen et al., 2022 | 5F8 | Z. parviglumis |
| 104 | Parviglumis_33 | Parviglumis33 | Chen et al., 2022 | 5F9 | Z. parviglumis |
| 105 | Parviglumis_34 | Parviglumis34 | Chen et al., 2022 | 5H10 | Z. parviglumis |
| 106 | Parviglumis_35 | Parviglumis35 | Chen et al., 2022 | 5H11 | Z. parviglumis |
| 107 | Parviglumis_36 | Parviglumis36 | Chen et al., 2022 | 5H12 | Z. parviglumis |
| 108 | Parviglumis_37 | Parviglumis37 | Chen et al., 2022 | 5H7 | Z. parviglumis |
| 109 | Parviglumis_38 | Parviglumis38 | Chen et al., 2022 | 5H8 | Z. parviglumis |
| 110 | Parviglumis_39 | Parviglumis39 | Chen et al., 2022 | 5H9 | Z. parviglumis |
| 111 | Parviglumis_40 | Parviglumis40 | Chen et al., 2022 | 6C10 | Z. parviglumis |
| 112 | Parviglumis_41 | Parviglumis41 | Chen et al., 2022 | 6C11 | Z. parviglumis |
| 113 | Parviglumis_42 | Parviglumis42 | Chen et al., 2022 | 6C12 | Z. parviglumis |
| 114 | Parviglumis_43 | Parviglumis43 | Chen et al., 2022 | 6D1 | Z. parviglumis |
| 115 | Parviglumis_44 | Parviglumis44 | Chen et al., 2022 | 6D10 | Z. parviglumis |
| 116 | Parviglumis_45 | Parviglumis45 | Chen et al., 2022 | 6D12 | Z. parviglumis |
| 117 | Parviglumis_46 | Parviglumis46 | Chen et al., 2022 | 6D2 | Z. parviglumis |
| 118 | Parviglumis_47 | Parviglumis47 | Chen et al., 2022 | 6D3 | Z. parviglumis |
| 119 | Parviglumis_48 | Parviglumis48 | Chen et al., 2022 | 6D4 | Z. parviglumis |
| 120 | Parviglumis_49 | Parviglumis49 | Chen et al., 2022 | 6D5 | Z. parviglumis |
| 121 | Parviglumis_50 | Parviglumis50 | Chen et al., 2022 | 6D6 | Z. parviglumis |
| 122 | Parviglumis_51 | Parviglumis51 | Chen et al., 2022 | 6D7 | Z. parviglumis |
| 123 | Parviglumis_52 | Parviglumis52 | Chen et al., 2022 | 6D8 | Z. parviglumis |
| 124 | Parviglumis_53 | Parviglumis53 | Chen et al., 2022 | 6D9 | Z. parviglumis |
| 125 | Parviglumis_54 | Parviglumis54 | Chen et al., 2022 | 6E2 | Z. parviglumis |
| 126 | Parviglumis_55 | Parviglumis55 | Chen et al., 2022 | 6E3 | Z. parviglumis |
| 127 | Parviglumis_56 | Parviglumis56 | Chen et al., 2022 | 6E4 | Z. parviglumis |
| 128 | Parviglumis_57 | Parviglumis57 | Chen et al., 2022 | 6E7 | Z. parviglumis |
| 129 | Parviglumis_58 | Parviglumis58 | Chen et al., 2022 | 6F11 | Z. parviglumis |
| 130 | Parviglumis_59 | Parviglumis59 | Chen et al., 2022 | 6F12 | Z. parviglumis |

| | | | | | |
|---|---|---|---|---|---|
| 131 | Parviglumis_60 | Parviglumis60 | Chen et al., 2022 | 6G11 | Z. parviglumis |
| 132 | Parviglumis_61 | Parviglumis61 | Chen et al., 2022 | 6G6 | Z. parviglumis |
| 133 | Parviglumis_62 | Parviglumis62 | Chen et al., 2022 | 6H1 | Z. parviglumis |
| 134 | Parviglumis_63 | Parviglumis63 | Chen et al., 2022 | 6H5 | Z. parviglumis |
| 135 | Parviglumis_64 | Parviglumis64 | Chen et al., 2022 | 6H6 | Z. parviglumis |
| 136 | Parviglumis_65 | Parviglumis65 | Chen et al., 2022 | 6H7 | Z. parviglumis |
| 137 | Parviglumis_66 | Parviglumis66 | Chen et al., 2022 | 6H8 | Z. parviglumis |
| 138 | Parviglumis_67 | Parviglumis67 | Chen et al., 2022 | 6H9 | Z. parviglumis |
| 139 | Parviglumis_68 | Parviglumis68 | Chia et al. 2012 | TIL01 | Z. parviglumis |
| 140 | Parviglumis_69 | Parviglumis69 | Chia et al. 2012 | TIL02 | Z. parviglumis |
| 141 | Parviglumis_70 | Parviglumis70 | Chia et al. 2012 | TIL03 | Z. parviglumis |
| 142 | Parviglumis_71 | Parviglumis71 | Chia et al. 2012 | TIL05 | Z. parviglumis |
| 143 | Parviglumis_72 | Parviglumis72 | Chia et al. 2012 | TIL06 | Z. parviglumis |
| 144 | Parviglumis_73 | Parviglumis73 | Chia et al. 2012 | TIL07 | Z. parviglumis |
| 145 | Parviglumis_74 | Parviglumis74 | Chia et al. 2012 | TIL08 | Z. parviglumis |
| 146 | Parviglumis_75 | Parviglumis75 | Chia et al. 2012 | TIL09 | Z. parviglumis |
| 147 | Parviglumis_76 | Parviglumis76 | Chia et al. 2012 | TIL10 | Z. parviglumis |
| 148 | Parviglumis_77 | Parviglumis77 | Chia et al. 2012 | TIL11 | Z. parviglumis |
| 149 | Parviglumis_78 | Parviglumis78 | Chia et al. 2012 | TIL12 | Z. parviglumis |
| 150 | Parviglumis_79 | Parviglumis79 | Chia et al. 2012 | TIL14 | Z. parviglumis |
| 151 | Parviglumis_80 | Parviglumis80 | Chia et al. 2012 | TIL15 | Z. parviglumis |
| 152 | Parviglumis_81 | Parviglumis81 | Chia et al. 2012 | TIL16 | Z. parviglumis |
| 153 | Parviglumis_82 | Parviglumis82 | Chia et al. 2012 | TIL17 | Z. parviglumis |
| 154 | Parviglumis_83 | Parviglumis83 | Chia et al. 2012 | TIL25 | Z. parviglumis |
| 155 | Parviglumis_84 | Parviglumis84 | Unterseer et al. 2014 | Teosinte | Z. parviglumis |
| 156 | Parviglumis_85 | Parviglumis85 | Wang et al. 2017 | parviglumis_2A | Z. parviglumis |
| 157 | Parviglumis_86 | Parviglumis86 | Wang et al. 2017 | parviglumis_2B | Z. parviglumis |
| 158 | Parviglumis_87 | Parviglumis87 | Wang et al. 2017 | parviglumis_2C | Z. parviglumis |
| 159 | Parviglumis_88 | Parviglumis88 | Wang et al. 2017 | parviglumis_2D | Z. parviglumis |
| 160 | Tripsacum | tripsacum | Chen et al., 2022 | tripsacum | Tripsacum dactyloides |
| 161 | Zmexicana_1 | Zmexicana1 | Chen et al., 2022 | 5A1 | Z. mexicana |
| 162 | Zmexicana_2 | Zmexicana2 | Chen et al., 2022 | 5A2 | Z. mexicana |
| 163 | Zmexicana_3 | Zmexicana3 | Chen et al., 2022 | 5A3 | Z. mexicana |

| | | | | | |
|---|---|---|---|---|---|
| 164 | Zmexicana_4 | Zmexicana4 | Chen et al., 2022 | 5A4 | Z. mexicana |
| 165 | Zmexicana_5 | Zmexicana5 | Chen et al., 2022 | 5A5 | Z. mexicana |
| 166 | Zmexicana_6 | Zmexicana6 | Chen et al., 2022 | 5A6 | Z. mexicana |
| 167 | Zmexicana_7 | Zmexicana7 | Chen et al., 2022 | 5A7 | Z. mexicana |
| 168 | Zmexicana_8 | Zmexicana8 | Chen et al., 2022 | 5B7 | Z. mexicana |
| 169 | Zmexicana_9 | Zmexicana9 | Chen et al., 2022 | 5B8 | Z. mexicana |
| 170 | Zmexicana_10 | Zmexicana10 | Chen et al., 2022 | 5B9 | Z. mexicana |
| 171 | Zmexicana_11 | Zmexicana11 | Chen et al., 2022 | 5D9 | Z. mexicana |
| 172 | Zmexicana_12 | Zmexicana12 | Chen et al., 2022 | 5E10 | Z. mexicana |
| 173 | Zmexicana_13 | Zmexicana13 | Chen et al., 2022 | 5E11 | Z. mexicana |
| 174 | Zmexicana_14 | Zmexicana14 | Chen et al., 2022 | 5E12 | Z. mexicana |
| 175 | Zmexicana_15 | Zmexicana15 | Chen et al., 2022 | 5E7 | Z. mexicana |
| 176 | Zmexicana_16 | Zmexicana16 | Chen et al., 2022 | 5E8 | Z. mexicana |
| 177 | Zmexicana_17 | Zmexicana17 | Chen et al., 2022 | 5E9 | Z. mexicana |
| 178 | Zmexicana_18 | Zmexicana18 | Chen et al., 2022 | 5F1 | Z. mexicana |
| 179 | Zmexicana_19 | Zmexicana19 | Chen et al., 2022 | 5F10 | Z. mexicana |
| 180 | Zmexicana_20 | Zmexicana20 | Chen et al., 2022 | 5F12 | Z. mexicana |
| 181 | Zmexicana_21 | Zmexicana21 | Chen et al., 2022 | 5F2 | Z. mexicana |
| 182 | Zmexicana_22 | Zmexicana22 | Chen et al., 2022 | 5F3 | Z. mexicana |
| 183 | Zmexicana_23 | Zmexicana23 | Chen et al., 2022 | 5F4 | Z. mexicana |
| 184 | Zmexicana_24 | Zmexicana24 | Chen et al., 2022 | 5F5 | Z. mexicana |
| 185 | Zmexicana_25 | Zmexicana25 | Chen et al., 2022 | 5F6 | Z. mexicana |
| 186 | Zmexicana_26 | Zmexicana26 | Chen et al., 2022 | 5F7 | Z. mexicana |
| 187 | Zmexicana_27 | Zmexicana27 | Chen et al., 2022 | 5G1 | Z. mexicana |
| 188 | Zmexicana_28 | Zmexicana28 | Chen et al., 2022 | 5G10 | Z. mexicana |
| 189 | Zmexicana_29 | Zmexicana29 | Chen et al., 2022 | 5G12 | Z. mexicana |
| 190 | Zmexicana_30 | Zmexicana30 | Chen et al., 2022 | 5G2 | Z. mexicana |
| 191 | Zmexicana_31 | Zmexicana31 | Chen et al., 2022 | 5G3 | Z. mexicana |
| 192 | Zmexicana_32 | Zmexicana32 | Chen et al., 2022 | 5G6 | Z. mexicana |
| 193 | Zmexicana_33 | Zmexicana33 | Chen et al., 2022 | 5G7 | Z. mexicana |
| 194 | Zmexicana_34 | Zmexicana34 | Chen et al., 2022 | 5G8 | Z. mexicana |
| 195 | Zmexicana_35 | Zmexicana35 | Chen et al., 2022 | 5G9 | Z. mexicana |
| 196 | Zmexicana_36 | Zmexicana36 | Chen et al., 2022 | 5H1 | Z. mexicana |

| | | | | | |
|---|---|---|---|---|---|
| 197 | Zmexicana_37 | Zmexicana37 | Chen et al., 2022 | 5H2 | Z. mexicana |
| 198 | Zmexicana_38 | Zmexicana38 | Chen et al., 2022 | 5H3 | Z. mexicana |
| 199 | Zmexicana_39 | Zmexicana39 | Chen et al., 2022 | 5H4 | Z. mexicana |
| 200 | Zmexicana_40 | Zmexicana40 | Chen et al., 2022 | 5H5 | Z. mexicana |
| 201 | Zmexicana_41 | Zmexicana41 | Chen et al., 2022 | 6A1 | Z. mexicana |
| 202 | Zmexicana_42 | Zmexicana42 | Chen et al., 2022 | 6A10 | Z. mexicana |
| 203 | Zmexicana_43 | Zmexicana43 | Chen et al., 2022 | 6A11 | Z. mexicana |
| 204 | Zmexicana_44 | Zmexicana44 | Chen et al., 2022 | 6A12 | Z. mexicana |
| 205 | Zmexicana_45 | Zmexicana45 | Chen et al., 2022 | 6A2 | Z. mexicana |
| 206 | Zmexicana_46 | Zmexicana46 | Chen et al., 2022 | 6A3 | Z. mexicana |
| 207 | Zmexicana_47 | Zmexicana47 | Chen et al., 2022 | 6A4 | Z. mexicana |
| 208 | Zmexicana_48 | Zmexicana48 | Chen et al., 2022 | 6A5 | Z. mexicana |
| 209 | Zmexicana_49 | Zmexicana49 | Chen et al., 2022 | 6A6 | Z. mexicana |
| 210 | Zmexicana_50 | Zmexicana50 | Chen et al., 2022 | 6A7 | Z. mexicana |
| 211 | Zmexicana_51 | Zmexicana51 | Chen et al., 2022 | 6A8 | Z. mexicana |
| 212 | Zmexicana_52 | Zmexicana52 | Chen et al., 2022 | 6A9 | Z. mexicana |
| 213 | Zmexicana_53 | Zmexicana53 | Chen et al., 2022 | 6B10 | Z. mexicana |
| 214 | Zmexicana_54 | Zmexicana54 | Chen et al., 2022 | 6B11 | Z. mexicana |
| 215 | Zmexicana_55 | Zmexicana55 | Chen et al., 2022 | 6B12 | Z. mexicana |
| 216 | Zmexicana_56 | Zmexicana56 | Chen et al., 2022 | 6B2 | Z. mexicana |
| 217 | Zmexicana_57 | Zmexicana57 | Chen et al., 2022 | 6B3 | Z. mexicana |
| 218 | Zmexicana_58 | Zmexicana58 | Chen et al., 2022 | 6B4 | Z. mexicana |
| 219 | Zmexicana_59 | Zmexicana59 | Chen et al., 2022 | 6B5 | Z. mexicana |
| 220 | Zmexicana_60 | Zmexicana60 | Chen et al., 2022 | 6B6 | Z. mexicana |
| 221 | Zmexicana_61 | Zmexicana61 | Chen et al., 2022 | 6B7 | Z. mexicana |
| 222 | Zmexicana_62 | Zmexicana62 | Chen et al., 2022 | 6B8 | Z. mexicana |
| 223 | Zmexicana_63 | Zmexicana63 | Chen et al., 2022 | 6B9 | Z. mexicana |
| 224 | Zmexicana_64 | Zmexicana64 | Chen et al., 2022 | 6C1 | Z. mexicana |
| 225 | Zmexicana_65 | Zmexicana65 | Chen et al., 2022 | 6C2 | Z. mexicana |
| 226 | Zmexicana_66 | Zmexicana66 | Chen et al., 2022 | 6C3 | Z. mexicana |
| 227 | Zmexicana_67 | Zmexicana67 | Chen et al., 2022 | 6C4 | Z. mexicana |
| 228 | Zmexicana_68 | Zmexicana68 | Chen et al., 2022 | 6C5 | Z. mexicana |
| 229 | Zmexicana_69 | Zmexicana69 | Chen et al., 2022 | 6C6 | Z. mexicana |

| | | | | | |
|---|---|---|---|---|---|
| 230 | Zmexicana_70 | Zmexicana70 | Chen et al., 2022 | 6C7 | Z. mexicana |
| 231 | Zmexicana_71 | Zmexicana71 | Chen et al., 2022 | 6C8 | Z. mexicana |
| 232 | Zmexicana_72 | Zmexicana72 | Chen et al., 2022 | 6C9 | Z. mexicana |
| 233 | Zmexicana_73 | Zmexicana73 | Chen et al., 2022 | 6D11 | Z. mexicana |
| 234 | Zmexicana_74 | Zmexicana74 | Chen et al., 2022 | 6E1 | Z. mexicana |
| 235 | Zmexicana_75 | Zmexicana75 | Chen et al., 2022 | 6E10 | Z. mexicana |
| 236 | Zmexicana_76 | Zmexicana76 | Chen et al., 2022 | 6E5 | Z. mexicana |
| 237 | Zmexicana_77 | Zmexicana77 | Chen et al., 2022 | 6E9 | Z. mexicana |
| 238 | Zmexicana_78 | Zmexicana78 | Chen et al., 2022 | 6H2 | Z. mexicana |
| 239 | Zmexicana_79 | Zmexicana79 | Chen et al., 2022 | 6H4 | Z. mexicana |
| 240 | SweetCorn_1 | SweetCorn1 | Grzybowski et al. 2023 | 4554_INBRED | Z. mays Sweet corn |
| 241 | SweetCorn_2 | SweetCorn2 | Grzybowski et al. 2023 | 80-2 | Z. mays Sweet corn |
| 242 | SweetCorn_3 | SweetCorn3 | Grzybowski et al. 2023 | C15 | Z. mays Sweet corn |
| 243 | SweetCorn_4 | SweetCorn4 | Grzybowski et al. 2023 | C42 | Z. mays Sweet corn |
| 244 | SweetCorn_5 | SweetCorn5 | Grzybowski et al. 2023 | C68 | Z. mays Sweet corn |
| 245 | SweetCorn_6 | SweetCorn6 | Qiu et al. 2021 | CA-4 | Z. mays Sweet corn |
| 246 | SweetCorn_7 | SweetCorn7 | Grzybowski et al. 2023 | CL17 | Z. mays Sweet corn |
| 247 | SweetCorn_8 | SweetCorn8 | Grzybowski et al. 2023 | CL27 | Z. mays Sweet corn |
| 248 | SweetCorn_9 | SweetCorn9 | Grzybowski et al. 2023 | CO245 | Z. mays Sweet corn |
| 249 | SweetCorn_10 | SweetCorn10 | Bukowski et al. 2018 | CO255 | Z. mays Sweet corn |
| 250 | SweetCorn_11 | SweetCorn11 | Bukowski et al. 2018 | EP1 | Z. mays Sweet corn |
| 251 | SweetCorn_12 | SweetCorn12 | Unterseer et al. 2014 | F2 | Z. mays Sweet corn |
| 252 | SweetCorn_13 | SweetCorn13 | Bukowski et al. 2018 | F7 | Z. mays Sweet corn |
| 253 | SweetCorn_14 | SweetCorn14 | Qiu et al. 2021 | FC46 | Z. mays Sweet corn |
| 254 | SweetCorn_15 | SweetCorn15 | Grzybowski et al. 2023 | G22_T122 | Z. mays Sweet corn |
| 255 | SweetCorn_16 | SweetCorn16 | Grzybowski et al. 2023 | G3_T5a | Z. mays Sweet corn |
| 256 | SweetCorn_17 | SweetCorn17 | Qiu et al. 2021 | IA2132 | Z. mays Sweet corn |
| 257 | SweetCorn_18 | SweetCorn18 | Grzybowski et al. 2023 | Ia453 | Z. mays Sweet corn |
| 258 | SweetCorn_19 | SweetCorn19 | Bukowski et al. 2018 | Ia5125 | Z. mays Sweet corn |
| 259 | SweetCorn_20 | SweetCorn20 | Grzybowski et al. 2023 | Ia5125B | Z. mays Sweet corn |
| 260 | SweetCorn_21 | SweetCorn21 | Bukowski et al. 2018 | Il101 | Z. mays Sweet corn |
| 261 | SweetCorn_22 | SweetCorn22 | Bukowski et al. 2018 | Il14H | Z. mays Sweet corn |
| 262 | SweetCorn_23 | SweetCorn23 | Grzybowski et al. 2023 | Il778d | Z. mays Sweet corn |

| | | | | | |
|---|---|---|---|---|---|
| 263 | SweetCorn_24 | SweetCorn24 | Grzybowski et al. 2023 | Il_101T | Z. mays Sweet corn |
| 264 | SweetCorn_25 | SweetCorn25 | Grzybowski et al. 2023 | NO._380 | Z. mays Sweet corn |
| 265 | SweetCorn_26 | SweetCorn26 | Bukowski et al. 2018 | P39 | Z. mays Sweet corn |
| 266 | SweetCorn_27 | SweetCorn27 | Grzybowski et al. 2023 | PHDD6 | Z. mays Sweet corn |
| 267 | SweetCorn_28 | SweetCorn28 | Grzybowski et al. 2023 | PHEM7 | Z. mays Sweet corn |
| 268 | SweetCorn_29 | SweetCorn29 | Grzybowski et al. 2023 | PHGG7 | Z. mays Sweet corn |
| 269 | SweetCorn_30 | SweetCorn30 | Grzybowski et al. 2023 | S_56 | Z. mays Sweet corn |
| 270 | SweetCorn_31 | SweetCorn31 | Grzybowski et al. 2023 | T146 | Z. mays Sweet corn |
| 271 | SweetCorn_32 | SweetCorn32 | Qiu et al. 2021 | T242 | Z. mays Sweet corn |
| 272 | SweetCorn_33 | SweetCorn33 | Grzybowski et al. 2023 | T9 | Z. mays Sweet corn |
| 273 | SweetCorn_34 | SweetCorn34 | Grzybowski et al. 2023 | U_123 | Z. mays Sweet corn |
| 274 | SweetCorn_35 | SweetCorn35 | Bukowski et al. 2018 | i1677a | Z. mays Sweet corn |
| 275 | Ztropical_1 | Ztropical1 | Grzybowski et al. 2023 | 4F-306_108 | Z. mays Tropical |
| 276 | Ztropical_2 | Ztropical2 | Qiu et al. 2021 | 4F-35_BK | Z. mays Tropical |
| 277 | Ztropical_3 | Ztropical3 | Qiu et al. 2021 | A272 | Z. mays Tropical |
| 278 | Ztropical_4 | Ztropical4 | Grzybowski et al. 2023 | A3G-3-3-1-313 | Z. mays Tropical |
| 279 | Ztropical_5 | Ztropical5 | Bukowski et al. 2018 | CML10 | Z. mays Tropical |
| 280 | Ztropical_6 | Ztropical6 | Bukowski et al. 2018 | CML11 | Z. mays Tropical |
| 281 | Ztropical_7 | Ztropical7 | Bukowski et al. 2018 | CML14 | Z. mays Tropical |
| 282 | Ztropical_8 | Ztropical8 | Bukowski et al. 2018 | CML157Q | Z. mays Tropical |
| 283 | Ztropical_9 | Ztropical9 | Bukowski et al. 2018 | CML158Q | Z. mays Tropical |
| 284 | Ztropical_10 | Ztropical10 | Bukowski et al. 2018 | CML238 | Z. mays Tropical |
| 285 | Ztropical_11 | Ztropical11 | Bukowski et al. 2018 | CML258 | Z. mays Tropical |
| 286 | Ztropical_12 | Ztropical12 | Bukowski et al. 2018 | CML261 | Z. mays Tropical |
| 287 | Ztropical_13 | Ztropical13 | Bukowski et al. 2018 | CML281 | Z. mays Tropical |
| 288 | Ztropical_14 | Ztropical14 | Bukowski et al. 2018 | CML311 | Z. mays Tropical |
| 289 | Ztropical_15 | Ztropical15 | Bukowski et al. 2018 | CML314 | Z. mays Tropical |
| 290 | Ztropical_16 | Ztropical16 | Bukowski et al. 2018 | CML321 | Z. mays Tropical |
| 291 | Ztropical_17 | Ztropical17 | Bukowski et al. 2018 | CML331 | Z. mays Tropical |
| 292 | Ztropical_18 | Ztropical18 | Bukowski et al. 2018 | CML332 | Z. mays Tropical |
| 293 | Ztropical_19 | Ztropical19 | Bukowski et al. 2018 | CML333 | Z. mays Tropical |
| 294 | Ztropical_20 | Ztropical20 | Bukowski et al. 2018 | CML341 | Z. mays Tropical |
| 295 | Ztropical_21 | Ztropical21 | Bukowski et al. 2018 | CML38 | Z. mays Tropical |

| | | | | | |
|---|---|---|---|---|---|
| 296 | Ztropical_22 | Ztropical22 | Bukowski et al. 2018 | CML45 | Z. mays Tropical |
| 297 | Ztropical_23 | Ztropical23 | Bukowski et al. 2018 | CML5 | Z. mays Tropical |
| 298 | Ztropical_24 | Ztropical24 | Bukowski et al. 2018 | CML61 | Z. mays Tropical |
| 299 | Ztropical_25 | Ztropical25 | Bukowski et al. 2018 | CML69 | Z. mays Tropical |
| 300 | Ztropical_26 | Ztropical26 | Bukowski et al. 2018 | CML_108 | Z. mays Tropical |
| 301 | Ztropical_27 | Ztropical27 | Bukowski et al. 2018 | CML_154Q | Z. mays Tropical |
| 302 | Ztropical_28 | Ztropical28 | Bukowski et al. 2018 | CML_218 | Z. mays Tropical |
| 303 | Ztropical_29 | Ztropical29 | Bukowski et al. 2018 | CML_220 | Z. mays Tropical |
| 304 | Ztropical_30 | Ztropical30 | Bukowski et al. 2018 | CML_228 | Z. mays Tropical |
| 305 | Ztropical_31 | Ztropical31 | Bukowski et al. 2018 | CML_247 | Z. mays Tropical |
| 306 | Ztropical_32 | Ztropical32 | Bukowski et al. 2018 | CML_254 | Z. mays Tropical |
| 307 | Ztropical_33 | Ztropical33 | Bukowski et al. 2018 | CML_264 | Z. mays Tropical |
| 308 | Ztropical_34 | Ztropical34 | Bukowski et al. 2018 | CML_277 | Z. mays Tropical |
| 309 | Ztropical_35 | Ztropical35 | Bukowski et al. 2018 | CML_287 | Z. mays Tropical |
| 310 | Ztropical_36 | Ztropical36 | Bukowski et al. 2018 | CML_322 | Z. mays Tropical |
| 311 | Ztropical_37 | Ztropical37 | Bukowski et al. 2018 | CML_323 | Z. mays Tropical |
| 312 | Ztropical_38 | Ztropical38 | Grzybowski et al. 2023 | CML_395 | Z. mays Tropical |
| 313 | Ztropical_39 | Ztropical39 | Bukowski et al. 2018 | CML_52 | Z. mays Tropical |
| 314 | Ztropical_40 | Ztropical40 | Bukowski et al. 2018 | CML_91 | Z. mays Tropical |
| 315 | Ztropical_41 | Ztropical41 | Qiu et al. 2021 | F2834T | Z. mays Tropical |
| 316 | Ztropical_42 | Ztropical42 | Grzybowski et al. 2023 | Hi28 | Z. mays Tropical |
| 317 | Ztropical_43 | Ztropical43 | Qiu et al. 2021 | Huanyao | Z. mays Tropical |
| 318 | Ztropical_44 | Ztropical44 | Grzybowski et al. 2023 | INBRED_100 | Z. mays Tropical |
| 319 | Ztropical_45 | Ztropical45 | Grzybowski et al. 2023 | INBRED_109 | Z. mays Tropical |
| 320 | Ztropical_46 | Ztropical46 | Grzybowski et al. 2023 | INBRED_2-687 | Z. mays Tropical |
| 321 | Ztropical_47 | Ztropical47 | Grzybowski et al. 2023 | INBRED_305 | Z. mays Tropical |
| 322 | Ztropical_48 | Ztropical48 | Grzybowski et al. 2023 | INBRED_309 | Z. mays Tropical |
| 323 | Ztropical_49 | Ztropical49 | Bukowski et al. 2018 | Ki11 | Z. mays Tropical |
| 324 | Ztropical_50 | Ztropical50 | Bukowski et al. 2018 | Ki14 | Z. mays Tropical |
| 325 | Ztropical_51 | Ztropical51 | Bukowski et al. 2018 | Ki2021 | Z. mays Tropical |
| 326 | Ztropical_52 | Ztropical52 | Bukowski et al. 2018 | Ki3 | Z. mays Tropical |
| 327 | Ztropical_53 | Ztropical53 | Bukowski et al. 2018 | Ki43 | Z. mays Tropical |
| 328 | Ztropical_54 | Ztropical54 | Bukowski et al. 2018 | Ki44 | Z. mays Tropical |

| | | | | | |
|---|---|---|---|---|---|
| 329 | Ztropical_55 | Ztropical55 | Wang et al. 2020 | MO18W | Z. mays Tropical |
| 330 | Ztropical_56 | Ztropical56 | Bukowski et al. 2018 | NC264 | Z. mays Tropical |
| 331 | Ztropical_57 | Ztropical57 | Bukowski et al. 2018 | NC296 | Z. mays Tropical |
| 332 | Ztropical_58 | Ztropical58 | Bukowski et al. 2018 | NC296A | Z. mays Tropical |
| 333 | Ztropical_59 | Ztropical59 | Bukowski et al. 2018 | NC298 | Z. mays Tropical |
| 334 | Ztropical_60 | Ztropical60 | Bukowski et al. 2018 | NC300 | Z. mays Tropical |
| 335 | Ztropical_61 | Ztropical61 | Bukowski et al. 2018 | NC302 | Z. mays Tropical |
| 336 | Ztropical_62 | Ztropical62 | Bukowski et al. 2018 | NC304 | Z. mays Tropical |
| 337 | Ztropical_63 | Ztropical63 | Bukowski et al. 2018 | NC320 | Z. mays Tropical |
| 338 | Ztropical_64 | Ztropical64 | Bukowski et al. 2018 | NC336 | Z. mays Tropical |
| 339 | Ztropical_65 | Ztropical65 | Bukowski et al. 2018 | NC338 | Z. mays Tropical |
| 340 | Ztropical_66 | Ztropical66 | Bukowski et al. 2018 | NC340 | Z. mays Tropical |
| 341 | Ztropical_67 | Ztropical67 | Bukowski et al. 2018 | NC346 | Z. mays Tropical |
| 342 | Ztropical_68 | Ztropical68 | Bukowski et al. 2018 | NC348 | Z. mays Tropical |
| 343 | Ztropical_69 | Ztropical69 | Bukowski et al. 2018 | NC350 | Z. mays Tropical |
| 344 | Ztropical_70 | Ztropical70 | Bukowski et al. 2018 | NC352 | Z. mays Tropical |
| 345 | Ztropical_71 | Ztropical71 | Bukowski et al. 2018 | NC354 | Z. mays Tropical |
| 346 | Ztropical_72 | Ztropical72 | Bukowski et al. 2018 | NC356 | Z. mays Tropical |
| 347 | Ztropical_73 | Ztropical73 | Bukowski et al. 2018 | NC358 | Z. mays Tropical |
| 348 | Ztropical_74 | Ztropical74 | Qiu et al. 2021 | NY6371 | Z. mays Tropical |
| 349 | Ztropical_75 | Ztropical75 | Bukowski et al. 2018 | Tx601 | Z. mays Tropical |
| 350 | Ztropical_76 | Ztropical76 | Bukowski et al. 2018 | Tzi10 | Z. mays Tropical |
| 351 | Ztropical_77 | Ztropical77 | Bukowski et al. 2018 | Tzi11 | Z. mays Tropical |
| 352 | Ztropical_78 | Ztropical78 | Bukowski et al. 2018 | Tzi18 | Z. mays Tropical |
| 353 | Ztropical_79 | Ztropical79 | Bukowski et al. 2018 | Tzi8 | Z. mays Tropical |
| 354 | Ztropical_80 | Ztropical80 | Bukowski et al. 2018 | Tzi9 | Z. mays Tropical |
| 355 | Ztropical_81 | Ztropical81 | Qiu et al. 2021 | W803G | Z. mays Tropical |
| 356 | Ztropical_82 | Ztropical82 | Bukowski et al. 2018 | WIL500 | Z. mays Tropical |
| 357 | Ztropical_83 | Ztropical83 | Grzybowski et al. 2023 | YANG | Z. mays Tropical |
| 358 | Ztropical_84 | Ztropical84 | Grzybowski et al. 2023 | YE-CHI-HUNG | Z. mays Tropical |
| 359 | Ztropical_85 | Ztropical85 | Grzybowski et al. 2023 | YELLOW_3-4 | Z. mays Tropical |
| 360 | Ztropical_86 | Ztropical86 | Grzybowski et al. 2023 | YE_4 | Z. mays Tropical |
| 361 | Europe_1 | Europe1 | Unterseer et al. 2014 | CH10 | Z. mays Europe |

| | | | | | |
|---|---|---|---|---|---|
| 362 | Europe_2 | Europe2 | Unterseer et al. 2014 | D06 | Z. mays Europe |
| 363 | Europe_3 | Europe3 | Unterseer et al. 2014 | D09 | Z. mays Europe |
| 364 | Europe_4 | Europe4 | Unterseer et al. 2014 | D152 | Z. mays Europe |
| 365 | Europe_5 | Europe5 | Unterseer et al. 2014 | DK105 | Z. mays Europe |
| 366 | Europe_6 | Europe6 | Unterseer et al. 2014 | EC169 | Z. mays Europe |
| 367 | Europe_7 | Europe7 | Unterseer et al. 2014 | EC49A | Z. mays Europe |
| 368 | Europe_8 | Europe8 | Unterseer et al. 2014 | EP44 | Z. mays Europe |
| 369 | Europe_9 | Europe9 | Unterseer et al. 2014 | EZ5 | Z. mays Europe |
| 370 | Europe_10 | Europe10 | Unterseer et al. 2014 | F03802 | Z. mays Europe |
| 371 | Europe_11 | Europe11 | Unterseer et al. 2014 | F252 | Z. mays Europe |
| 372 | Europe_12 | Europe12 | Unterseer et al. 2014 | F283 | Z. mays Europe |
| 373 | Europe_13 | Europe13 | Unterseer et al. 2014 | F353 | Z. mays Europe |
| 374 | Europe_14 | Europe14 | Unterseer et al. 2014 | F618 | Z. mays Europe |
| 375 | Europe_15 | Europe15 | Unterseer et al. 2014 | F64 | Z. mays Europe |
| 376 | Europe_16 | Europe16 | Unterseer et al. 2014 | F98902 | Z. mays Europe |
| 377 | Europe_17 | Europe17 | Unterseer et al. 2014 | FF0721H-7 | Z. mays Europe |
| 378 | Europe_18 | Europe18 | Unterseer et al. 2014 | Lo11 | Z. mays Europe |
| 379 | Europe_19 | Europe19 | Grzybowski et al. 2023 | S018693 | Z. mays Europe |
| 380 | Europe_20 | Europe20 | Grzybowski et al. 2023 | S03198 | Z. mays Europe |
| 381 | Europe_21 | Europe21 | Grzybowski et al. 2023 | S160 | Z. mays Europe |
| 382 | Europe_22 | Europe22 | Grzybowski et al. 2023 | S245 | Z. mays Europe |
| 383 | Europe_23 | Europe23 | Grzybowski et al. 2023 | S25 | Z. mays Europe |
| 384 | Europe_24 | Europe24 | Grzybowski et al. 2023 | S311 | Z. mays Europe |
| 385 | Europe_25 | Europe25 | Grzybowski et al. 2023 | S336A | Z. mays Europe |
| 386 | Europe_26 | Europe26 | Grzybowski et al. 2023 | S50676 | Z. mays Europe |
| 387 | Europe_27 | Europe27 | Grzybowski et al. 2023 | S61328 | Z. mays Europe |
| 388 | Europe_28 | Europe28 | Grzybowski et al. 2023 | S68911 | Z. mays Europe |
| 389 | Europe_29 | Europe29 | Grzybowski et al. 2023 | S84854 | Z. mays Europe |
| 390 | Europe_30 | Europe30 | Unterseer et al. 2014 | UH006 | Z. mays Europe |
| 391 | Europe_31 | Europe31 | Unterseer et al. 2014 | UH007 | Z. mays Europe |
| 392 | Europe_32 | Europe32 | Unterseer et al. 2014 | UH009 | Z. mays Europe |
| 393 | Europe_33 | Europe33 | Unterseer et al. 2014 | UH250 | Z. mays Europe |
| 394 | Europe_34 | Europe34 | Unterseer et al. 2014 | UH304 | Z. mays Europe |
