## Supplemental1 Dataset S2 for "Fifteenth century CE Bolivian maize reveals genetic affinities with ancient Peruvian maize"

**Supplementary Dataset 2.** Known SNPs related traits.

| SNPs | aBM_frq | ancient_frq | Type | Gene_name | Biologoical_process | SNPs and traits |
|---|---|---|---|---|---|---|
| 3_5721506 | G:1 A:0 | G:-nan A:-nan | TYPE=splice_donor_variant&intron_variant;EFFECT=HIGH;GENEMODEL=Zm00001eb120960 | bhlh70 | GO:0006355: regulation of transcription, DNA-templated | NA |
| 3_13134097 | G:1 A:0 | G:-nan A:-nan | TYPE=stop_gained;EFFECT=HIGH;GENEMODEL=Zm00001eb123120 | NA | GO:0006470: protein dephosphorylation | transcript SNP chromosome position structure trait 3:13183499; chr3; 13132965; Flanking region internode length below ear |
| 4_76831415 | A:1 G:0 | A:-nan G:-nan | TYPE=stop_lost;EFFECT=HIGH;GENEMODEL=Zm00001eb178600 | NA | GO:0006508: proteolysis | transcript SNP chromosome position structure trait 4:73852529; chr4; 76830220; Flanking region; nodes above ear |
| 5_128622059 | A:1 T:0 | A:-nan T:-nan | TYPE=stop_gained;EFFECT=HIGH;GENEMODEL=Zm00001eb236660 | NA | NA | NA |
