## Supplemental Dataset S3 for "Fifteenth century CE Bolivian maize reveals genetic affinities with ancient Peruvian maize"

**Supplementary Dataset 3.** SNPs informcation from SNPVersity.

| Chromosome | Location | Ref | Alt | MQ | Type | Effect | Genemodel |
|---|---|---|---|---|---|---|---|
| chr1 | 100470406 | T | A | MQ=. | TYPE=5_prime_UTR_variant | EFFECT=MODIFIER | GENEMODEL=Zm00001eb024430 |
| chr1 | 120053462 | G | A | MQ=. | TYPE=3_prime_UTR_variant | EFFECT=MODIFIER | GENEMODEL=Zm00001eb026360 |
| chr1 | 198168469 | C | T | MQ=. | TYPE=5_prime_UTR_variant | EFFECT=MODIFIER | GENEMODEL=Zm00001eb036550 |
| chr1 | 21693299 | T | G | MQ=. | TYPE=3_prime_UTR_variant | EFFECT=MODIFIER | GENEMODEL=Zm00001eb007310 |
| chr1 | 232277209 | C | T | MQ=. | TYPE=3_prime_UTR_variant | EFFECT=MODIFIER | GENEMODEL=Zm00001eb044430 |
| chr1 | 232277218 | C | T | MQ=. | TYPE=3_prime_UTR_variant | EFFECT=MODIFIER | GENEMODEL=Zm00001eb044430 |
| chr1 | 281885099 | C | T | MQ=. | TYPE=3_prime_UTR_variant | EFFECT=MODIFIER | GENEMODEL=Zm00001eb056940 |
| chr1 | 281885122 | G | A | MQ=. | TYPE=3_prime_UTR_variant | EFFECT=MODIFIER | GENEMODEL=Zm00001eb056940 |
| chr1 | 281885124 | A | C | MQ=. | TYPE=3_prime_UTR_variant | EFFECT=MODIFIER | GENEMODEL=Zm00001eb056940 |
| chr1 | 294107836 | T | C | MQ=. | TYPE=5_prime_UTR_variant | EFFECT=MODIFIER | GENEMODEL=Zm00001eb060520 |
| chr1 | 301697138 | C | T | MQ=. | TYPE=5_prime_UTR_variant | EFFECT=MODIFIER | GENEMODEL=Zm00001eb063130 |
| chr1 | 47420482 | A | G | MQ=. | TYPE=5_prime_UTR_variant | EFFECT=MODIFIER | GENEMODEL=Zm00001eb014140 |
| chr2 | 12612806 | A | C | MQ=. | TYPE=5_prime_UTR_variant | EFFECT=MODIFIER | GENEMODEL=Zm00001eb071790 |
| chr2 | 155177009 | C | G | MQ=. | TYPE=5_prime_UTR_variant | EFFECT=MODIFIER | GENEMODEL=Zm00001eb093800 |
| chr2 | 208927157 | A | G | MQ=. | TYPE=5_prime_UTR_variant | EFFECT=MODIFIER | GENEMODEL=Zm00001eb105710 |
| chr2 | 214458566 | T | C | MQ=. | TYPE=3_prime_UTR_variant | EFFECT=MODIFIER | GENEMODEL=Zm00001eb107820 |
| chr2 | 225583353 | G | C | MQ=. | TYPE=3_prime_UTR_variant | EFFECT=MODIFIER | GENEMODEL=Zm00001eb112130 |
| chr2 | 232834414 | A | C | MQ=. | TYPE=5_prime_UTR_variant | EFFECT=MODIFIER | GENEMODEL=Zm00001eb114270 |
| chr2 | 240323011 | G | A | MQ=. | TYPE=5_prime_UTR_variant | EFFECT=MODIFIER | GENEMODEL=Zm00001eb117240 |
| chr2 | 241646841 | C | A | MQ=. | TYPE=5_prime_UTR_variant,3_prime_UTR_variant | EFFECT=MODIFIER,MODIFIER | GENEMODEL=Zm00001eb117810,Zm00001eb117800 |
| chr2 | 29376800 | C | A | MQ=. | TYPE=3_prime_UTR_variant | EFFECT=MODIFIER | GENEMODEL=Zm00001eb077140 |
| chr2 | 39867034 | A | C | MQ=. | TYPE=5_prime_UTR_variant | EFFECT=MODIFIER | GENEMODEL=Zm00001eb079800 |
| chr2 | 51645650 | T | G | MQ=. | TYPE=3_prime_UTR_variant | EFFECT=MODIFIER | GENEMODEL=Zm00001eb082590 |
| chr2 | 9338102 | G | C | MQ=. | TYPE=3_prime_UTR_variant | EFFECT=MODIFIER | GENEMODEL=Zm00001eb070240 |
| chr3 | 151302151 | G | A | MQ=. | TYPE=5_prime_UTR_variant | EFFECT=MODIFIER | GENEMODEL=Zm00001eb140980 |
| chr3 | 212283868 | A | C | MQ=. | TYPE=5_prime_UTR_variant,3_prime_UTR_variant | EFFECT=MODIFIER,MODIFIER | GENEMODEL=Zm00001eb156100,Zm00001eb156110 |
| chr3 | 216097910 | G | A | MQ=. | TYPE=5_prime_UTR_variant | EFFECT=MODIFIER | GENEMODEL=Zm00001eb157320 |
| chr3 | 225047941 | T | C | MQ=. | TYPE=3_prime_UTR_variant | EFFECT=MODIFIER | GENEMODEL=Zm00001eb160100 |
| chr3 | 225346147 | T | C | MQ=. | TYPE=3_prime_UTR_variant | EFFECT=MODIFIER | GENEMODEL=Zm00001eb160200 |
| chr3 | 24515288 | G | C | MQ=. | TYPE=5_prime_UTR_variant | EFFECT=MODIFIER | GENEMODEL=Zm00001eb125660 |
| chr3 | 8064756 | C | T | MQ=. | TYPE=3_prime_UTR_variant | EFFECT=MODIFIER | GENEMODEL=Zm00001eb121570 |
| chr4 | 161508528 | T | C | MQ=. | TYPE=5_prime_UTR_variant | EFFECT=MODIFIER | GENEMODEL=Zm00001eb187340 |
| chr4 | 16159721 | A | G | MQ=. | TYPE=5_prime_UTR_variant | EFFECT=MODIFIER | GENEMODEL=Zm00001eb168890 |
| chr4 | 186157135 | G | A | MQ=. | TYPE=3_prime_UTR_variant | EFFECT=MODIFIER | GENEMODEL=Zm00001eb194260 |
| chr4 | 204933584 | C | T | MQ=. | TYPE=3_prime_UTR_variant | EFFECT=MODIFIER | GENEMODEL=Zm00001eb199740 |
| chr4 | 204934807 | A | T | MQ=. | TYPE=3_prime_UTR_variant | EFFECT=MODIFIER | GENEMODEL=Zm00001eb199740 |
| chr4 | 222041395 | T | C | MQ=. | TYPE=5_prime_UTR_variant | EFFECT=MODIFIER | GENEMODEL=Zm00001eb202560 |
| chr4 | 234903022 | T | G | MQ=. | TYPE=5_prime_UTR_premature_start_codon_gain_variant,5_prime_UTR_variant,3_prime_UTR_variant | EFFECT=LOW,MODIFIER,MODIFIER | GENEMODEL=Zm00001eb204470,Zm00001eb204470,Zm00001eb204480 |
| chr4 | 238320402 | C | A | MQ=. | TYPE=5_prime_UTR_variant | EFFECT=MODIFIER | GENEMODEL=Zm00001eb205130 |
| chr4 | 248765537 | G | A | MQ=. | TYPE=5_prime_UTR_variant | EFFECT=MODIFIER | GENEMODEL=Zm00001eb209550 |
| chr4 | 33315514 | C | T | MQ=. | TYPE=5_prime_UTR_variant | EFFECT=MODIFIER | GENEMODEL=Zm00001eb172250 |
| chr4 | 74539366 | C | A | MQ=. | TYPE=5_prime_UTR_variant | EFFECT=MODIFIER | GENEMODEL=Zm00001eb178330 |
| chr4 | 80602381 | G | A | MQ=. | TYPE=5_prime_UTR_variant | EFFECT=MODIFIER | GENEMODEL=Zm00001eb179000 |
| chr4 | 8324819 | T | C | MQ=. | TYPE=3_prime_UTR_variant | EFFECT=MODIFIER | GENEMODEL=Zm00001eb167520 |
| chr5 | 10577906 | A | G | MQ=. | TYPE=3_prime_UTR_variant | EFFECT=MODIFIER | GENEMODEL=Zm00001eb216010 |
| chr5 | 1206936 | T | G | MQ=. | TYPE=5_prime_UTR_variant | EFFECT=MODIFIER | GENEMODEL=Zm00001eb210550 |
| chr5 | 12277409 | G | C | MQ=. | TYPE=5_prime_UTR_variant | EFFECT=MODIFIER | GENEMODEL=Zm00001eb216590 |
| chr5 | 12556547 | T | A | MQ=. | TYPE=5_prime_UTR_variant | EFFECT=MODIFIER | GENEMODEL=Zm00001eb216680 |
| chr5 | 157165607 | T | C | MQ=. | TYPE=5_prime_UTR_variant | EFFECT=MODIFIER | GENEMODEL=Zm00001eb239920 |
| chr5 | 176024793 | C | T | MQ=. | TYPE=5_prime_UTR_variant | EFFECT=MODIFIER | GENEMODEL=Zm00001eb243620 |
| chr5 | 182418121 | T | A | MQ=. | TYPE=3_prime_UTR_variant | EFFECT=MODIFIER | GENEMODEL=Zm00001eb245580 |
| chr5 | 198246143 | G | T | MQ=. | TYPE=5_prime_UTR_variant | EFFECT=MODIFIER | GENEMODEL=Zm00001eb250130 |
| chr5 | 198341182 | G | A | MQ=. | TYPE=5_prime_UTR_variant | EFFECT=MODIFIER | GENEMODEL=Zm00001eb250170 |
| chr5 | 70626783 | C | G | MQ=. | TYPE=3_prime_UTR_variant | EFFECT=MODIFIER | GENEMODEL=Zm00001eb229700 |
| chr6 | 110407148 | A | G | MQ=. | TYPE=5_prime_UTR_variant | EFFECT=MODIFIER | GENEMODEL=Zm00001eb276280 |
| chr6 | 147316776 | T | C | MQ=. | TYPE=5_prime_UTR_variant | EFFECT=MODIFIER | GENEMODEL=Zm00001eb285170 |
| chr6 | 153729777 | A | T | MQ=. | TYPE=3_prime_UTR_variant | EFFECT=MODIFIER | GENEMODEL=Zm00001eb286890 |
| chr6 | 162632708 | G | A | MQ=. | TYPE=3_prime_UTR_variant | EFFECT=MODIFIER | GENEMODEL=Zm00001eb289570 |
| chr6 | 177562480 | A | C | MQ=. | TYPE=5_prime_UTR_variant | EFFECT=MODIFIER | GENEMODEL=Zm00001eb296760 |
| chr6 | 46828966 | T | C | MQ=. | TYPE=5_prime_UTR_variant,3_prime_UTR_variant | EFFECT=MODIFIER,MODIFIER | GENEMODEL=Zm00001eb266960,Zm00001eb266970 |
| chr6 | 6532633 | C | T | MQ=. | TYPE=3_prime_UTR_variant | EFFECT=MODIFIER | GENEMODEL=Zm00001eb260040 |
| chr7 | 169286701 | T | G | MQ=. | TYPE=3_prime_UTR_variant | EFFECT=MODIFIER | GENEMODEL=Zm00001eb325340 |
| chr7 | 180327845 | G | A | MQ=. | TYPE=3_prime_UTR_variant | EFFECT=MODIFIER | GENEMODEL=Zm00001eb329850 |
| chr7 | 25747316 | T | C | MQ=. | TYPE=5_prime_UTR_variant | EFFECT=MODIFIER | GENEMODEL=Zm00001eb304360 |
| chr7 | 6339053 | T | G | MQ=. | TYPE=3_prime_UTR_variant | EFFECT=MODIFIER | GENEMODEL=Zm00001eb300250 |
| chr7 | 87611506 | G | A | MQ=. | TYPE=5_prime_UTR_variant | EFFECT=MODIFIER | GENEMODEL=Zm00001eb309370 |
| chr8 | 149732409 | A | C | MQ=. | TYPE=5_prime_UTR_variant | EFFECT=MODIFIER | GENEMODEL=Zm00001eb358930 |
| chr8 | 153975834 | A | G | MQ=. | TYPE=3_prime_UTR_variant | EFFECT=MODIFIER | GENEMODEL=Zm00001eb360090 |
| chr8 | 166563940 | C | A | MQ=. | TYPE=5_prime_UTR_variant | EFFECT=MODIFIER | GENEMODEL=Zm00001eb363850 |
| chr8 | 166603677 | T | C | MQ=. | TYPE=5_prime_UTR_variant | EFFECT=MODIFIER | GENEMODEL=Zm00001eb363870 |
| chr8 | 173191480 | A | G | MQ=. | TYPE=5_prime_UTR_variant | EFFECT=MODIFIER | GENEMODEL=Zm00001eb366800 |
| chr8 | 175400583 | T | A | MQ=. | TYPE=3_prime_UTR_variant | EFFECT=MODIFIER | GENEMODEL=Zm00001eb367930 |
| chr8 | 175486254 | T | G | MQ=. | TYPE=5_prime_UTR_variant | EFFECT=MODIFIER | GENEMODEL=Zm00001eb368010 |
| chr8 | 175486277 | T | C | MQ=. | TYPE=5_prime_UTR_variant | EFFECT=MODIFIER | GENEMODEL=Zm00001eb368010 |
| chr8 | 179273127 | T | C | MQ=. | TYPE=5_prime_UTR_variant | EFFECT=MODIFIER | GENEMODEL=Zm00001eb370130 |
| chr8 | 8181863 | G | A | MQ=. | TYPE=5_prime_UTR_variant | EFFECT=MODIFIER | GENEMODEL=Zm00001eb334280 |
| chr9 | 10690588 | T | C | MQ=. | TYPE=5_prime_UTR_variant | EFFECT=MODIFIER | GENEMODEL=Zm00001eb373560 |
| chr9 | 114213148 | A | T | MQ=. | TYPE=3_prime_UTR_variant | EFFECT=MODIFIER | GENEMODEL=Zm00001eb389810 |
| chr9 | 15610991 | G | C | MQ=. | TYPE=5_prime_UTR_variant | EFFECT=MODIFIER | GENEMODEL=Zm00001eb374940 |
| chr9 | 46090129 | G | T | MQ=. | TYPE=3_prime_UTR_variant | EFFECT=MODIFIER | GENEMODEL=Zm00001eb381380 |
| chr9 | 67326318 | G | A | MQ=. | TYPE=5_prime_UTR_premature_start_codon_gain_variant,5_prime_UTR_variant | EFFECT=LOW,MODIFIER | GENEMODEL=Zm00001eb383260,Zm00001eb383260 |
| chr10 | 116200880 | C | A | MQ=. | TYPE=5_prime_UTR_variant | EFFECT=MODIFIER | GENEMODEL=Zm00001eb421830 |
| chr10 | 148023740 | G | A | MQ=. | TYPE=3_prime_UTR_variant | EFFECT=MODIFIER | GENEMODEL=Zm00001eb432340 |
| chr10 | 87628027 | G | A | MQ=. | TYPE=5_prime_UTR_variant | EFFECT=MODIFIER | GENEMODEL=Zm00001eb417310 |
