## Supplemental Tables for "Fifteenth century CE Bolivian maize reveals genetic affinities with ancient Peruvian maize"

**Table S1. Geographical information of the archaeological maize samples used in this study.**

| ID | Country | Age (BP) | Location | Latitude | Longitude | Altitude (m above sea level) |
|---|---|---|---|---|---|---|
| Arica4 | Chile | 990 +/- 30 | Arica, Chile, coastal | 18.47S | 70.32W | coastal |
| Arica5 | Chile | 780 +/- 30 | Arica, Chile, coastal | 18.47S | 70.32W | coastal |
| Z2 | Brazil | 570 +- 60 | Brazil:Peruacu Valley, Boquete cave | 15S | 44W | 700 |
| Z6 | Brazil | 630+- 60 | Brazil:Peruacu Valley, Lapa de Hora | 15S | 44W | 700 |
| Z61 | Peru | 800 +/- 30 | Peru:Ancash | 8.531S | 78.341W | 200 |
| Z64 | Peru | 630 +/- 30 | Peru:Ancash | 9.064S | 77.564W | 3990 |
| Z65 | Peru | 970 +/- 30 | Peru:Inca | 14.433S | 75.342W | 220 |
| Z66 | Argentina | 1010 +/- 30 | Argentina:Catamerca | 27.342S | 66.550W | 2465 |
| Z67 | Argentina | 100 +/- 30 | Argentina:Jujuy | 23.240S | 66.211W | 3700 |
| EG84 | Honduras | 1870 – 1740 | El Gigante rock shelter | 14.22N | 88.06W | NA |
| EG85 | Honduras | 2300 – 2070 | El Gigante rock shelter | 14.22N | 88.06W | NA |
| EG90 | Honduras | 2300 – 2120 | El Gigante rock shelter | 14.22N | 88.06W | NA |
| Tehua162 | Mexico | 5310 | Mexico: Tehuacan, Puebla | 18.46N | 97.39W | NA |
| SM3 | Mexico | 4190 +/- 30 | Mexico: San Marcos cave, Tehuacan Valley | 17.48N | 97.03W | NA |
| SM5 | Mexico | 4190 +/- 30 | Mexico: San Marcos cave, Tehuacan Valley | 17.48N | 97.03W | NA |
| SM10 | Mexico | 4240 +/- 30 | Mexico: San Marcos cave, Tehuacan Valley | 17.48N | 97.03W | NA |

* The latitude and longitude information for SM3 and SM10 were determined based on the Tehuacan Valley location as provided in 'Myxomycetes associated with dryland ecosystems of the Tehuacán-Cuicatlán Valley Biosphere Reserve, Mexico."

**Table S2. Paleogenetic characterization of archaeological Bolivian maize sequence samples with six libraries.**

| Sample | QC-passed reads | Total number of read mapped | Read mapped (%) | Mean Coverage | Std Coverage |
|---|---|---|---|---|---|
| aBMComb_rm5nt.rmDR.bam | 299093147 | 33846381 | 11.32% | 0.7667X | 3.027X |
| Bolivian_Maize_1_ATCACG_L001_rm5nt.rmDR.bam | 56128831 | 6402435 | 11.41% | 0.1434X | 0.6598X |
| Bolivian_Maize_2_CGATGT_L001_rm5nt.rmDR.bam | 67115071 | 8050825 | 12.00% | 0.1815X | 0.7718X |
| Bolivian_Maize_3_TTAGGC_L001_rm5nt.rmDR.bam | 53043360 | 5998148 | 11.31% | 0.1399X | 0.6726X |
| Bolivian_Maize_4_TGACCA_L001_rm5nt.rmDR.bam | 10941693 | 1579877 | 14.44% | 0.0371X | 0.2633X |
| Bolivian_Maize_5_ACAGTG_L001_rm5nt.rmDR.bam | 10694963 | 1652909 | 15.46% | 0.0361X | 0.2482X |
| Bolivian_Maize_6_GCCAAT_L001_rm5nt.rmDR.bam | 101169229 | 10162187 | 10.04% | 0.2287X | 0.9215X |

**Table S3. The percentage of genomic sites covered at variable depths in the archaeological Bolivian maize (aBM) sample.** (aBMComb_rm5nt.rmDR.bam)

| Depth | aBM_Combbam |
|---|---|
| 1X | 27.75% |
| 2X | 14.14% |
| 3X | 8.27% |
| 4X | 5.18% |
| 5X | 3.41% |
| 6X | 2.35% |
| 7X | 1.7% |
| 8X | 1.28% |
| 9X | 1% |
| >=10X | 9.78% |
